# Modular and programmable RNA sensing using ADAR editing in living cells

**DOI:** 10.1101/2022.01.28.478207

**Authors:** K. Eerik Kaseniit, Noa Katz, Natalie S. Kolber, Connor C. Call, Diego L. Wengier, Will B. Cody, Elizabeth S. Sattely, Xiaojing J. Gao

## Abstract

With the increasing availability of single-cell transcriptomes, RNA signatures offer a promising basis for targeting living cells. Molecular RNA sensors would enable the study of and therapeutic interventions for specific cell types/stats in diverse contexts, particularly in human patients and non-model organisms. Here we describe a modular and programmable design for live **<u>R</u>**NA sensing using **<u>a</u>**denosine **<u>d</u>**eaminases **<u>a</u>**cting on **<u>R</u>**NA (RADAR). We validated and then expanded our basic design, characterized its performance, and thoroughly analyzed its compatibility with the human/mouse transcriptomes. We also identified strategies to further boost output levels and improve the dynamic range. We show that RADAR is programmable and modular, and uniquely enables compact AND logic. In addition to being quantitative, RADAR can distinguish disease-relevant point mutations. Finally, we demonstrate that RADAR is a self-contained system with the potential to function in diverse organisms.

## Introduction

Identifying signature transcripts for cell types or states of interest has now become routine thanks to great progress in bulk and single-cell transcriptome analysis^1^. However, it is often laborious or very challenging to genetically access those same states, for example to turn on a reporter or other gene of interest only in cells expressing a given transcript. The ability to express arbitrary proteins in response to arbitrary RNAs would allow for the functional dissection of neural circuits, the detection and ablation of cancer cells, and myriad additional applications in both research and therapeutics that are enabled by the precise manipulation of cell subpopulations.

Currently, cell types and states are routinely manipulated using promoter engineering, by selecting promoter sequences that mirror the expression of marker transcripts expressed in cell states of interest and placing genes of interest under the control of the engineered promoter^2^. However, such engineering is not guaranteed to succeed, as it is by necessity bespoke, not compact as promoters can be several kilobases in size, and prone to unpredictable behavior due to unknown transcription factor binding sites that are inadvertently included or omitted from the synthetic promoter.

Alternatively, one could create a molecular sensor that detects the RNA directly. The ideal sensor design would be (i) modular, with the inputs and outputs easily swapped with predictable outcomes, (ii) programmable for detecting all transcripts in a genome, (iii) amenable to combining several sensors in a parallel (multiplexed) way and capable of compact logic in a small nucleotide footprint, (iv) able to discriminate between minor nucleotide changes, such as single nucleotide variants, (v) useful in model and non-model organisms.

Due to the predictability and programmability of base pairing, the sensor is likely best made using RNA itself. So far, progress on this front has focused on manipulating specific, functional RNA structures within sensor RNAs, such that an input RNA would disrupt or reconstitute a functional RNA structure such as a guide RNA (gRNA) ^3–6^, or internal ribosome entry site (IRES)^7^ to enable or disable downstream protein translation. However, no existing design using these functional RNA sequences has met all the five criteria outlined above, largely due to two major challenges. First, these designs incorporate functional RNA sequences that must form specific secondary and tertiary structures beyond simple base-pairing in order to function appropriately and effectively, and this can make sensor RNA design difficult or unpredictable, reducing modularity and programmability. Second, double-stranded RNA (dsRNA) structures form as intermediates during sensing, but this can trigger unwanted responses as eukaryotic cells have many mechanisms to detect and respond to unexpected dsRNA^8^, adding another layer of complexity and unpredictability not accounted for in the existing sensors. We reason that instead of engineering a mechanism to circumvent the second aspect and further complicating sensor design, it could be leveraged as a feature instead.

In human cells, there are three main responses to dsRNAs: immune responses, RNA interference (RNAi), and editing by adenosine deaminases acting on RNA (ADAR)^8^. The immune response is not suitable for modular and programmable sensors as the endogenous transcriptional response would have to be diverted or modified. RNAi has been leveraged to create modular sensors for miRNAs^9–11^, demonstrating the feasibility of embracing endogenous dsRNA-engaging pathways as sensing mechanisms. However, RNAi-based sensing can directly only generate an output in the absence of an input (NOT logic), and it is not trivial to extend it to the sensing of mRNAs, so far only demonstrated in cell lysates^11^ and only using extensive chemical modification in living cells^10^. In contrast, we hypothesized that the third response, ADAR editing, provides a promising mechanism for designing RNA sensors using transcribed unmodified mRNAs. ADARs are present in all metazoa, and humans have three ADARs: ADAR1 (with two major isoforms), ADAR2, and the catalytically inactive inhibitory ADAR3^12^. All ADARs have non-specific dsRNA-binding domains, and ADAR1/2 have functional catalytic domains that convert adenosines (A) to inosines (I) within dsRNA. ADAR editing stands out compared to the other responses by the highly specific changes (A→I) generated in response to an input (dsRNA) largely agnostic of the sequence identity, which is important for modularity and programmability. ADAR editing also tends to bias the cellular response away from the two other pathways, as A-to-I edited dsRNA stretches are poor substrates for the immune receptors and RNAi^13,14^, further reducing unintended interactions between the sensor and the cellular context. In addition to the intrinsic benefits of ADAR, our engineering efforts are facilitated by the rich knowledge from an active field that has investigated the basic biology of ADAR^15–19^ and is applying it to the editing of disease-causing transcripts in gene therapies^20–23^.

Here we describe a design for **<u>R</u>**NA sensing using **<u>ADAR</u>** (RADAR). We show that this design meets the five criteria described above: modularity, programmability, multiplexing and logic capabilities, sensitivity to nucleotide-level variations, and potential to function in many organisms.

## Results

We termed our sensor architecture based on ADAR editing RADAR. Central to RADAR is a “sensor mRNA,” which contains two coding sequences (CDS), a marker (mCherry) and an output (EGFP), separated by a “sensor sequence” containing an in-frame UAG stop codon (**Figure 1a**). The sensor sequence is designed to be reverse complementary to the target RNA of interest (referred to as “trigger” henceforth), but with an editing-enhancing C:A mismatch (see below) at the UAG stop, such that the trigger RNA must contain a CCA subsequence. The stop codon prevents the downstream output EGFP coding sequence from being translated so that only mCherry is expressed. However, in the presence of the trigger RNA, a double-stranded RNA (dsRNA) stretch is formed around the stop codon by the sensor and trigger RNAs, which results in recruitment of ADAR enzymes. ADAR enzymes will edit the adenosine (A) in the UAG to an inosine (I), with efficiency increased by a C:A mismatch, resulting in UIG in the senor mRNA. While the C:A mismatch in this pairing is optimal, it is also possible to use 5’ GCA 3’, 5’ UCA 3’, and 5’ CAA 3’, as these sequences have also been shown to effectively edit a 5’ UAG 3’ stop to UIG^21^. UIG is read as UGG, encoding tryptophan, by the translational machinery, enabling the translation of the downstream CDS so that both mCherry and EGFP are expressed. “Self-cleaving” 2A sequences^24^ are introduced to insulate the sensor sequence from the flanking CDSs to avoid the induction of degradation, aggregates, or other undesired side effects by the peptide encoded in the sensor sequence.

**Figure 1.**
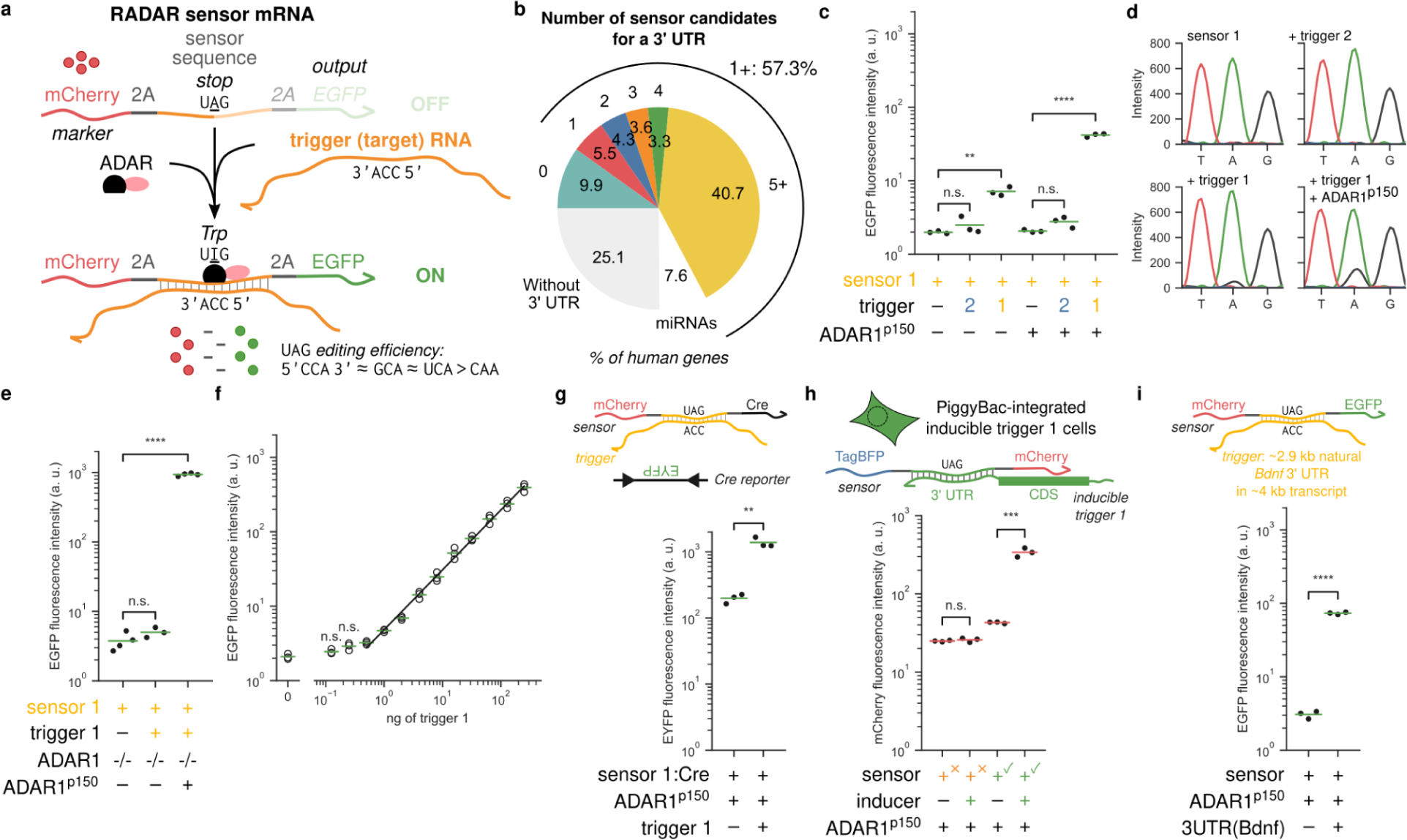
Modular live RNA sensing using ADAR editing. **a** The RADAR mechanism works via dsRNA-mediated A-to-I editing by ADAR, where a trigger RNA forms a dsRNA complex with a sensor mRNA, allowing a stop codon in the sensor to be edited away. mCherry is constitutively expressed as a marker, while the EGFP output is only expressed after the trigger RNA has bound the sensor sequence, ADAR recruited to the resulting dsRNA, and the UAG stop codon edited to UIG, which is read as encoding tryptophan. 2A sequences insulate the sensor sequence from the marker and output proteins, enabling modularity. UAG can be complemented by several sequences, with CCA, GCA and UCA resulting in similar editing efficiency, while CAA:UAG results in lower efficiency. **b** The majority of human genes have RADAR-compatible trigger subsequences in 3’ UTRs, with 41% of genes having over five candidate sensors. miRNAs can be detected with specialized techniques and were thus not analyzed. **c** RADAR functions in human cells in a sequence-specific manner and is enhanced by ADAR1^p150^ over-expression. Sensor 1 detects trigger 1, but not the non-matching trigger 2. **d** Sanger sequencing validating trigger-dependent editing and its enhancement by ADAR1^p150^ overexpression. A-to-I editing is evidenced by the presence of G signal (black) in the A position of Sanger traces. **e** RADAR functionality is ablated in ADAR1-deficient cells, and can be rescued with ADAR1^p150^ over-expression. **f** RADAR is quantitative over two logs of the amount of transfected trigger DNA. **g** RADAR output is modular, e.g., one can use the Cre recombinase as an alternative output. The reporter is turned on by Cre-mediated inversion of a EYFP CDS. **h** RADAR detects a genomically integrated trigger. The trigger-expressing transgene is under the control of a CMV-tetO promoter, induced by the addition of doxycycline. **i** RADAR detects a subsequence within a natural 3’ UTR. In this and subsequent figures, unless otherwise noted, we report the mean output fluorescence intensity in cells gated for high transfection efficiency (relative within the given experiment). Each dot represents one biological replicate and the horizontal lines indicate the mean of data in each group. Significance is determined by Bonferroni-corrected two-tailed Student t-test.

The 3’ untranslated regions (UTRs) of mRNAs are an attractive target for finding suitable trigger sequences over other regions of an mRNA due to three reasons: first, they are present in all forms of processed mRNA; second, they do not encounter translating ribosomes which may affect or be affected by the dsRNA intermediates formed; and third, any unintentional edits that occur in the 3’ UTR of a trigger mRNA are less likely to be detrimental compared to those in the CDS. We therefore analyzed all genes with annotated 3’ UTRs in the human genome in search of trigger sequences compatible with the RADAR design. Candidates must have a 5’ CCA 3’ sequence (which is paired with the 5’ UAG 3’ in the sensor) flanked by a sufficiently long stretch of sequence; the reverse complement of the trigger sequence should not create an in-frame stop codon in the sensor. Furthermore, the trigger sequences should be unique (assessed by mapping to the genome), and the sensor sequence easily synthesizable (lacking homopolymer runs and with 35-80% GC content). We identified that 57.3% of genes have a trigger candidate for at least one reported 3’ UTR variant (**Figure 1b**). Similar results were obtained for the mouse transcriptome (**Supplemental Figure 1c**).

To facilitate a fast design-build-test cycle, we first verified that RADAR works for a synthetic (*de novo* designed), plasmid-expressed trigger mRNA, T1. In the prototype sensor for T1, S1, a red mCherry fluorescent protein was constitutively expressed as the first CDS, and EGFP as the second (output) (**Figure 1a**). The initial optimization we performed included the validation that the 2A sequences improve the consistency of sensor behavior across different sensor sequences^24^, and that trigger-independent sensor output is minimized through the choice of the constitutive promoter driving sensor expression (**Supplemental Figure 1d**). We were able to establish a sensor specifically triggered by the T1 sequence embedded in the 3’ UTR of a cotransfected gene, as we saw that fluorescence output was increased only in the presence of the correct trigger (**Figure 1c**). As a more direct readout compared to the fluorescent output, we also validated A-to-I editing via Sanger sequencing of the sensor cDNA (**Figure 1d**). To validate that the sensor operates through the intended mechanism, in addition to the low endogenous expression of ADAR1 in HEK293 cells, we overexpressed several forms of ADAR, which indeed increased the sensor output (**Figure 1c, 1d, Supplemental Figure 1e**). We observed the best performance when overexpressing ADAR1^p150^ (**Supplemental Figure 1e**), the larger ADAR1 isoform that is interferon-induced and contains a Z-DNA/Z-RNA binding domain as well as a nuclear export signal in addition to the nuclear localization signal shared with the ADAR1^p110^ isoform^12^. Conversely, sensor output was ablated in ADAR1 knockout cells and rescued upon ADAR1^p150^ cotransfection (**Figure 1e**). We also characterized the monotonic relation between RADAR output and the amount of ADAR (**Supplemental Figure 1f**).

Because differences across cell types and states are often not only qualitative, but quantitative in mRNA expression levels, we investigated how RADAR responds to varying amounts of trigger. We observed that S1 output is linearly correlated with the amount of transfected T1 plasmid over two orders of magnitude (**Figure 1f**).

We next investigated whether RADAR output could be modularly switched, by changing the output to the Cre recombinase, a particularly useful tool in neurobiology^2^ that can catalyze the inversion of a reporter gene. RADAR→Cre is not only functional, but also enables output amplification, as we observed a greater signal in the presence of an input compared to when the fluorescent protein is expressed directly (**Figure 1g**). The increase in signal is at the cost of increased baseline final output (EYFP) expression, as a small baseline expression of Cre from the sensor is likely also amplified.

We then set to validate the functionality of RADAR in scenarios closer to the eventual use cases. First, in addition to transiently delivered trigger plasmids, we investigated whether we could sense genomically integrated trigger transcripts. We used piggyBac integration to generate a polyclonal cell line where the T1 sequence was placed in the 3’ UTR of a doxycycline-inducible EGFP coding sequence. We also designed a new sensor mRNA, swapping the marker from mCherry to TagBFP2 and the output from EGFP to mCherry, keeping other parts the same. We saw increased output fluorescence in response to doxycycline-induced trigger expression as expected (**Figure 1h**). The increased baseline signal of the correct sensor compared to the non-matching sensor might be explained by leaky EGFP expression in the uninduced case, as we observe significant EGFP expression in the uninduced case (**Supplemental Figure 2a-c**). With this EGFP-encoding trigger we were also able to investigate whether the dsRNA formed during sensing negatively impacts the sensed transcript. If the sensor decreased the trigger mRNA stability, lower transcript and protein levels would be observed. However, we did not observe any significant changes to the expression of EGFP (**Supplemental Figure 1g**).

After testing a synthetic trigger we next investigated whether RADAR would detect a trigger sequence in a more complex and natural context. We designed a sensor for a subsequence within the 2.9 kb 3’ UTR of the mouse *Bdnf* gene encoding the brain-derived neurotrophic factor, using the murine natural context in an orthogonal organism, human cells. When we expressed this natural murine 3’ UTR in human cells, we observed substantial increase of the sensor output compared to no trigger expression (**Figure 1i**), on par with our initial results with our prototype synthetic trigger and sensor pair.

While the majority of human genes have at least one trigger candidate, we investigated whether the repertoire of triggers could further be expanded. Particularly, one of the main reasons for a sequence not being a suitable trigger is the presence of in-frame stop codons in the reverse complement, which would result in a second, unedited stop codon in the sensor sequence. Two approaches could result in utilizing sequences that are less likely to contain in-frame stops in the reverse complement. First, we took advantage of the 3’ UTR of *Bdnf* to assess RADAR performance with different lengths of dsRNA, as a shorter sensor sequence is less likely to contain an in-frame stop codon. We observed that the performance of RADAR was largely maintained when the dsRNA sensor sequence is shortened to 72 bp or lengthened to 153 bp (**Figure 2a**). However, the trigger-induced output was abolished in a 36 bp sensor. Second, motivated by work showing that ADAR enzymes are tolerant of or even benefited by disruptions in the target dsRNA^25^, we hypothesized that we could rescue the signal from the non-functional 36 bp sensor if we introduced an additional 54 bp sequence complementary to another part of the long *Bdnf* 3’ UTR, so that a 90 bp discontiguous dsRNA would be formed between the sensor and trigger. A similar strategy of split complementarity has recently been described to benefit RNA editing with endogenous ADARs ^23^. A suitable short, 36 bp sensor sequence can be more readily identified in a transcript compared to a longer sequence, and nearly all transcripts have at least one 54 bp stretch without an in-frame stop in the reverse complement. We observed that this sensor for a “split” trigger was functional, (**Figure 2b**), albeit with slightly reduced activation compared to the contiguous 90 bp sensor. Finally, since not all genes have 3’ UTRs that are amenable to sensing, we also investigated whether the CDS of an mRNA could be a functional trigger to further expand RADAR’ s range. To test this, we used our inducible trigger-integrated cell line, which contained the coding sequence for EGFP. The EGFP CDS sensor (S_EGFP_) showed a significant but modest change in output in response to trigger induction (**Figure 2c**). The same subsequence of the EGFP CDS was able to more strongly trigger the sensor when placed in the 3’ UTR (**Supplemental Figure 2d**), consistent with our hypothesis that there may be interference of the dsRNA formation or editing by the translational machinery, which had motivated our initial targeting of 3’ UTRs rather than CDSs. In contrast to the 3’ UTR sensor, the CDS sensor did have a small negative effect on the EGFP fluorescence (**Supplemental Figure 2e**), potentially due to disrupting translation or other natural transcript processing. In summary, sensing CDS sequences can be used as an alternative when 3’ UTRs are not available for sensing. Combining the ability to shorten the sensor, using a split trigger, and sensing CDSs in addition to 3’ UTRs, we estimate that 85% of human genes have at least one candidate trigger sequence compatible with RADAR (**Figure 2d**); similar results were obtained for the mouse transcriptome (**Supplemental Figure 2f**). Notably, the largest group of human genes without trigger candidates are small nucleolar RNA genes, which make up 21% of such genes.

**Figure 2.**
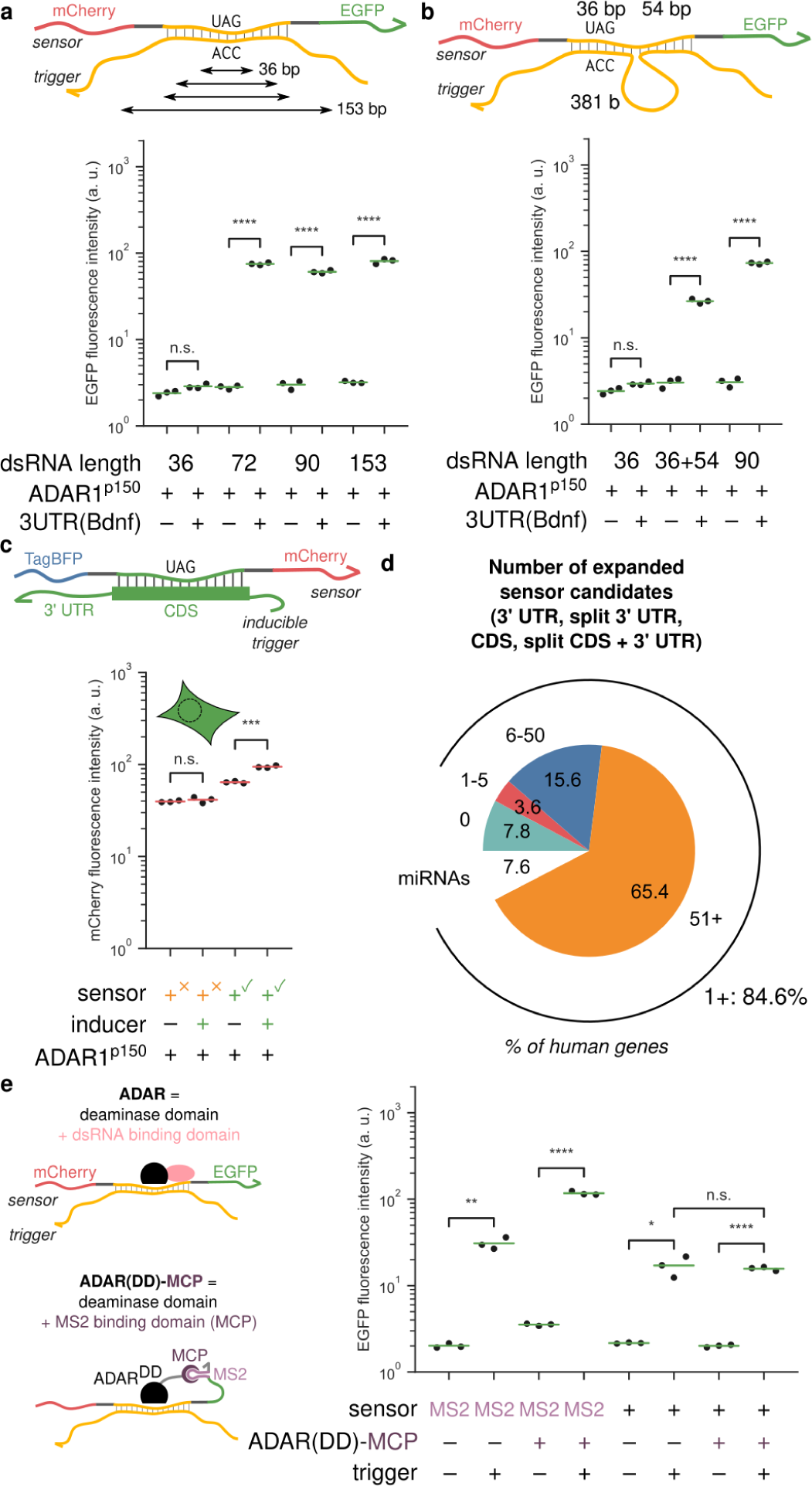
Expanding the range of RADAR and enhancing its output dynamic range. **a** The length of the dsRNA stretch formed between the sensor and trigger can be varied down to at least 72 bp or up to at least 153 bp without affecting sensor performance. **b** The sensor can be designed to form dsRNA with two “split” segments on the trigger sequence, allowing more flexibility in trigger choice. **c** The sensor detects a trigger sequence within a CDS of a gene. **d** The range of genes RADAR could target is expanded when sensor sequences can be shortened, a split trigger can be used, or if CDSs can be a source of triggers. **e** Enhancing RADAR using an engineered ADAR that only binds the sensor mRNA via MS2-MCP interactions. The engineered chimeric ADAR does not enhance editing of the sensor in the absence of the MS2 sequence.

While we consistently leveraged ADAR over-expression to demonstrate robust performance of RADAR, it may restrict its application, because overexpressed ADAR might increase the editing of endogenous dsRNA structures and be detrimental to certain cell types or settings. For example, ADAR overexpression was observed to significantly associate with increased size, metastasis and stage of gastric cancer tumors^26^. To ameliorate this potential problem, we engineered orthogonal recruitment of ADAR to the sensor mRNA, rather than relying on the wild-type ADAR binding to the sensor-trigger dsRNA with little sequence specificity. We used a fusion protein of the MS2 RNA motif-binding protein MCP to a hyperactive-editing variant of the deaminase domain of ADAR2 (“ADAR(DD)-MCP”)^20,27^. We expect ADAR(DD)-MCP to only specifically edit RNAs with the MS2 sequence, since a major source of affinity to arbitrary dsRNA has been removed and the deaminase domain on its own has stronger sequence biases than the dsRNA binding domains^28^. The best results were achieved when we placed three copies of the MS2 sequence into the 3’ UTR of our prototype S1 sensor, where we observed a greater output dynamic range for this sensor in the presence of ADAR(DD)-MCP compared to without any type of ADAR over-expression, and achieved a similar dynamic range when ADAR1^p150^ was over-expressed as before (**Figure 2e, Supplemental Figure 2g**). Crucially, ADAR(DD)-MCP co-expression did not alter the dynamic range of a sensor without MS2 sequences (**Figure 2e**), suggesting that this engineered “orthogonal” ADAR is unlikely to edit other endogenous dsRNA structures.

Having optimized and characterized RADAR, we next investigated its unique features and potential applications. We posited that RADAR could be used to classify cells, so tested this concept by mixing our T1-integrated cell line (**Figure 1g**) with the parental cell line and using the corresponding S1 (3’ UTR) or S_EGFP_ (EGFP CDS) sensors to detect cells expressing the trigger. In an approximately 1:1 mixture of EGFP^-^:EGFP^+^ cells, we first filtered for highly transfected cells based on the constitutive TagBFP2 expression from the sensor. Next we varied the mCherry output fluorescence threshold at which a cell was considered “positive” to establish the tradeoff between false and true positive rates. Finally, high or low EGFP fluorescence was to determine whether the classifier determination was a true or false positive, and in a receiver operating characteristic (ROC) analysis, the 3’ UTR sensor achieved an average area under the curve (AUC) of 0.85, the CDS sensor an AUC of 0.58, and an unrelated sensor an AUC of 0.53 (**Figure 3a**). The 3’ UTR sensor therefore performs well in the classification task, and even the suboptimal CDS sensor exhibits better-than-random performance.

**Figure 3.**
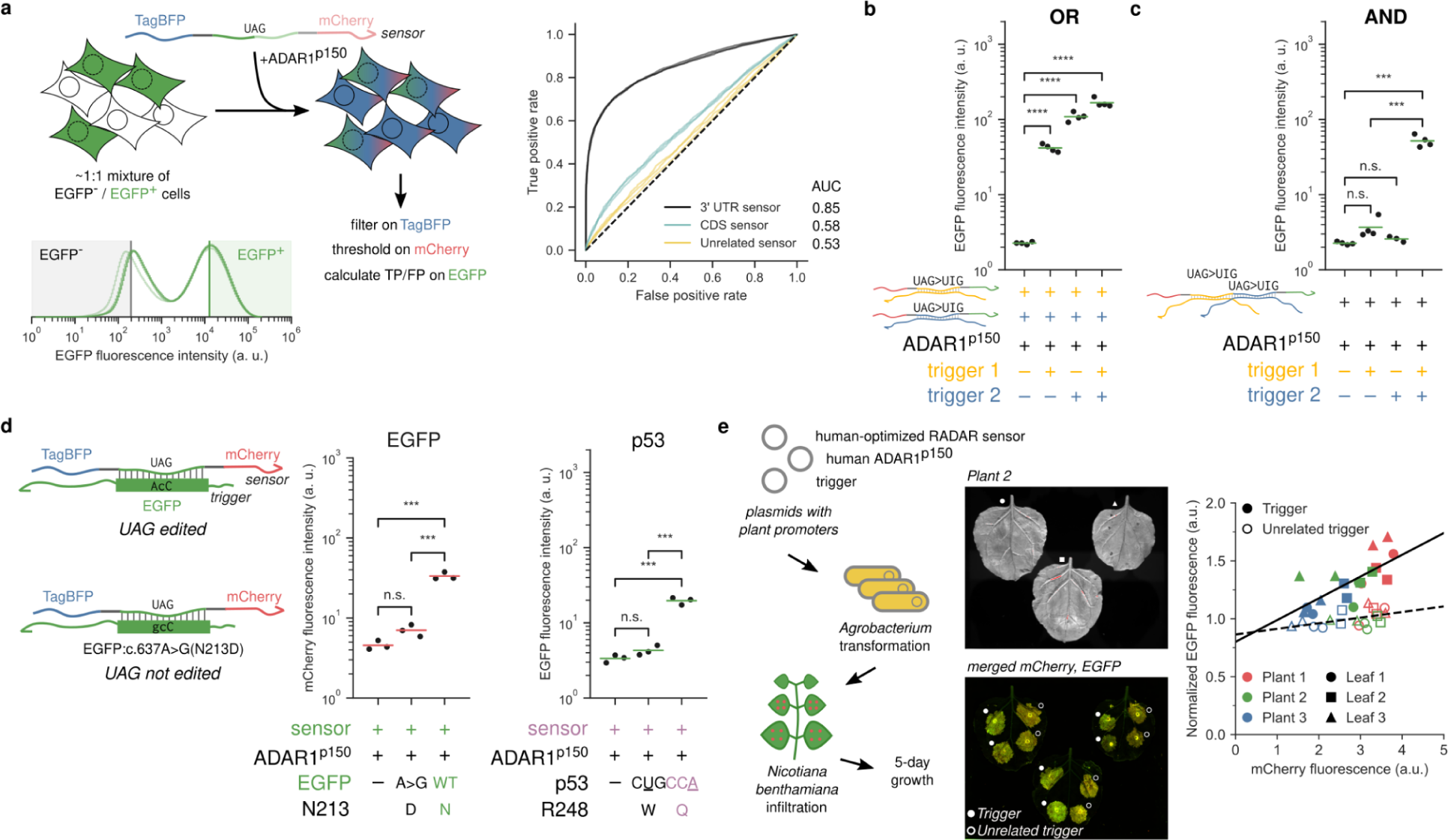
Unique features and potential applications of RADAR. **a** RADAR classifies cells based on the expression of a transgene. **b** OR logic (multiplexing) is achieved by the co-delivery of two sensors. **c** AND logic is achieved by placing two sensor sequences in series such that stop codons in both sequences have to be edited for expression of the output CDS in the sensor. **d** RADAR detects or distinguishes between point mutations. **e** RADAR works in a heterologous context in plants. Three leaves in three *Nicotiana benthamiana* plants were infiltrated with *Agrobacterium* separately transformed with RADAR components (sensor and human ADAR1^p150^), and a matching or unrelated trigger, with two infiltrations of either trigger per leaf. Images from a representative plant. Quantification based on averages of average fluorescence intensities from six rectangular regions per infiltration spot.

A single-input classifier may suffice in some, but not all scenarios. However, the RADAR architecture lends itself easily to OR-type logic (multiplexing), as two sensors with different sensor sequences but the same output can simply be co-delivered, as well as AND-type logic, as two different sensor sequences can be concatenated within one sensor mRNA, resulting in two sequential in-frame stop codons that both have to be edited in order for there to be translation of the downstream output coding sequence. We validated the performance of both gates (**Figure 3b, 3c**). These logic gates, especially AND, can enable more sophisticated targeting of cell sub-types and -states than single-input sensors prototyped above.

ADAR is highly sensitive to the base identities surrounding the edited adenosine on both strands of the dsRNA substrate^21^. Therefore, RADAR is uniquely suitable for the sensitive distinction of certain single-base variants. As a proof-of-principle, we introduced a point mutation into EGFP (A>G change at position 637 modifying CCA to CCG), a change reported to abrogate ADAR editing when opposite the UAG stop codon in the sensor. As expected, the CDS sensor we previously characterized responds only to the wild type but not to the mutant EGFP (**Figure 3d**). To demonstrate the biomedical potential of such allelic distinctions, we then created a sensor to distinguish between two common oncogenic mutations of *TP53*, R248W (CCG>CUG, predicted to not trigger RADAR) and R248Q (CCG>CCA, predicted to trigger RADAR). While these two mutations alter the same residue, it has been reported that the cells harboring the R248Q variant were more invasive than those with the R248W variant^29^. As expected, the sensor responds to the R248Q but not the R248W variant when the mutant cDNA was co-transfected (**Figure 3d**). We estimate that approximately 5% of known or likely pathogenic variants reported in ClinVar can potentially be distinguished from their wild type alleles using RADAR.

Finally, considering the reliance of RADAR only on the presence of complementary mRNA and the ADAR, we hypothesized that it could operate as a self-contained module. To test this, we assessed RADAR in plants which do not have endogenous ADAR expression. We used *Nicotiana benthamiana* as a model system. Without any optimization other than switching to plant promoters, we infiltrated *N. benthamiana* leaves with spots of *Agrobacterium* strains carrying plasmids encoding the S1 sensor, the T1 trigger, and ADAR. We observed elevated fluorescent output in response to the matching T1 trigger compared to an non-matching trigger (**Figure 3e**). This indicates that RADAR still operates as intended in a heterologous context, and is therefore potentially useful in a wide range of species, even those lacking native ADAR machinery.

## Discussion

In this study we demonstrated leveraging ADAR editing to create a modular molecular device capable of sensing near-arbitrary RNAs. RADAR met the criteria of modularity in terms of both inputs and outputs, an expansive range of targets, capability of multiplexing (OR logic) and compact AND logic, as well as sensitivity to small nucleotide changes. For applications where the potential for off-target dsRNA editing due to ADAR over-expression is undesirable, we demonstrated the functionality of a sensor-recruited engineered ADAR comprising the ADAR deaminase domain and a MS2 RNA binding domain. In addition, since the only non-sensor component required is ADAR activity, we showed that RADAR could work across kingdoms.

We experimentally investigated several strategies to expand the range of RADAR, and thoroughly analyzed RADAR’s compatibility with the human/mouse transcriptomes. We envision that the range of RADAR can be further expanded if necessary. For example, a problematic in-frame stop codon may be amenable to removal using a single-base mismatch that removes the stop codon in the sensor, particularly if the nuisance stop codon is far from the edited UAG. A second strategy could be to replace the UAG:CCA/GCA/UCA/CAA pairing with a less efficient yet still substantially edited pairing, such as UAG:ACA^21^. Third, an expression-correlated RNA could be used instead as a surrogate for the RNA of interest; two correlated RNAs could be combined using an AND gate for increased specificity. This bespoke engineering can be done rapidly by simply swapping out the short sensor sequence in a sensor plasmid using easily synthesized oligonucleotides.

Since RADAR is based on mRNAs and all components (sensor and ADAR or ADAR(DD)-MCP) can be delivered via mRNA, the wide range of existing RNA synthetic biology tools^30^ could be combined with RADAR to produce highly sophisticated cell classifiers, research tools, and smart dynamic therapies. For example, RADAR sensors could be used to track cells as they become infected with a virus, as they transition from normal to pre-cancerous to cancerous to metastatic, or as they become senescent to study these processes, or to create smart therapies that can dynamically detect these state changes, and stop or even reverse the processes driving them. By integrating multiple inputs, RADAR can enable high specificity and low off-target effects of the downstream interventions. It is especially suitable for increasing the specificity of RNA-based vaccination and gene therapies, the power of which was recently demonstrated during the pandemic.

We envision that RADAR could be particularly useful in non-model organisms and human patients where germline transgenesis is very limited or not feasible. As RADAR can be delivered on viral or other genetic vectors and achieve cell type-specific expression, it removes the need for promoter identification, which has remained a major hurdle in onboarding new organisms for bioengineering or genetics-driven research.

## Acknowledgments

This work was funded by NIH (4R00EB027723-02; X.J.G), Seed Grant from Brain Research Foundation (X.J.G), NARSAD Young Investigator Grant from Brain & Behavior Research Foundation (X.J.G), Stanford Bio-X Interdisciplinary Graduate Fellowship (K.E.K), Fulbright Foundation (N.K.), NSF GRFP (N.S.K), Stanford ChEM-H CBI training program (N.S.K), EDGE Doctoral Fellowship Program (N.S.K). Noa Katz is an Awardee of the Weizmann Institute of Science – Israel National Postdoctoral Award Program for Advancing Women in Science. We thank the Gao lab members for their feedback. We thank prof. Liqun Luo for gifts of Cre related plasmids, prof. Billy Li for ADAR plasmids and the ADAR1 knockout cell line, and prof. Robert Singer for the ADAR(DD)-MCP plasmid.

## Author contributions

K.E.K, N.K. N.S.K, and X.J.G designed the study. K.E.K, N.K. N.S.K, and C.C.C performed and analyzed most of the experiments, with support from D.L.W, W.B.C, and E.S.S for the plant experiments. K.E.K. performed bioinformatic analysis. K.E.K. and X.J.G. wrote the manuscript with input from all authors.

## Materials and methods

### Plasmid generation

Plasmids were generated using standard molecular cloning practices, including InFusion of linearized plasmids and PCR fragments and restriction-ligation of linearized fragments and annealed phosphorylated oligonucleotides. A complete list of plasmids and associated maps is found in **Supplemental Table 1**. Plasmids are available upon request from the corresponding author. Human ADAR plasmids as well as the ADAR1 knockout cell line were generously provided by prof. Billy Li. Cre and Cre reporter plasmids were kindly gifted by prof. Liqun Luo. pUBC_stdMCP_serinemod_E488QADAR_p2A_yGFP (“ADAR(DD)-MCP”) was a gift from Robert Singer (Addgene plasmid # 154787; http://n2t.net/addgene:154787; RRID:Addgene_154787).

### Tissue culture

Flp-In T-REx Human Embryonic Kidney (HEK) 293 cells were purchased from Thermo Scientific (catalog #R78007). Cells were cultured in a humidity-controlled incubator under standard culture conditions (37°C with 5% CO2) in Dulbecco’ s Modified Eagle Medium, supplemented with 10% fetal bovine serum (Fisher Scientific catalog #FB12999102), 1 mM sodium pyruvate (EMD Millipore catalog # TMS-005-C), 1x penicillin-streptomycin (Genesee catalog #25-512), 2 mM L-glutamine (Genesee catalog #25-509) and 1x MEM non-essential amino acids (Genesee catalog #25-536). To induce expression of certain constructs, 100 ng/mL of doxycycline was added at the time of transfection. The inducible-trigger containing cell line was generated by transfecting the construct in a PiggyBac backbone along with PiggyBac integrase (4:1), with 50 ug/mL hygromycin added for selection when reseeding into 10 cm dishes two days after transfection.

### Transient transfections

HEK 293T cells were cultured in either 24-well or 96-well tissue culture-treated plates under standard culture conditions. When cells were 70-90% confluent, the cells were transiently transfected with plasmid constructs using the jetOPTIMUS® DNA transfection Reagent (Polyplus catalog # 117-15), as per manufacturer’ s instructions using 0.375 uL of reagent per 50 uL of jetOPTIMUS buffer for 500 ng total DNA transfections in the 24-well format and 0.13 uL of reagent per 12.5 uL of buffer for 130 ng total DNA in the 96-well format. All transfections are detailed in **Supplemental Table 2**.

### Flow cytometry and data analysis

Cells were harvested approximately 48 hours after transfection by trypsinization and resuspended in flow buffer (HBSS + 2.5 mg/mL bovine serum albumin). Post 40 um straining, cells were analyzed by flow cytometry (Biorad ZE5 Cell Analyzer), and data was processed with the cytoflow Python package. An overview of gating and analysis is given in **Supplemental Figure 1a**.

### Statistical analysis

Values are reported as the means from at least three biological replicates, representative of two independent biological experiments. For experiments comparing two groups, a Bonferroni-corrected two-tailed Student t-test was used to assess significance.

### Data availability

All reported data are available on request from the corresponding author.

## Figures

**Supplemental Figure 1.**
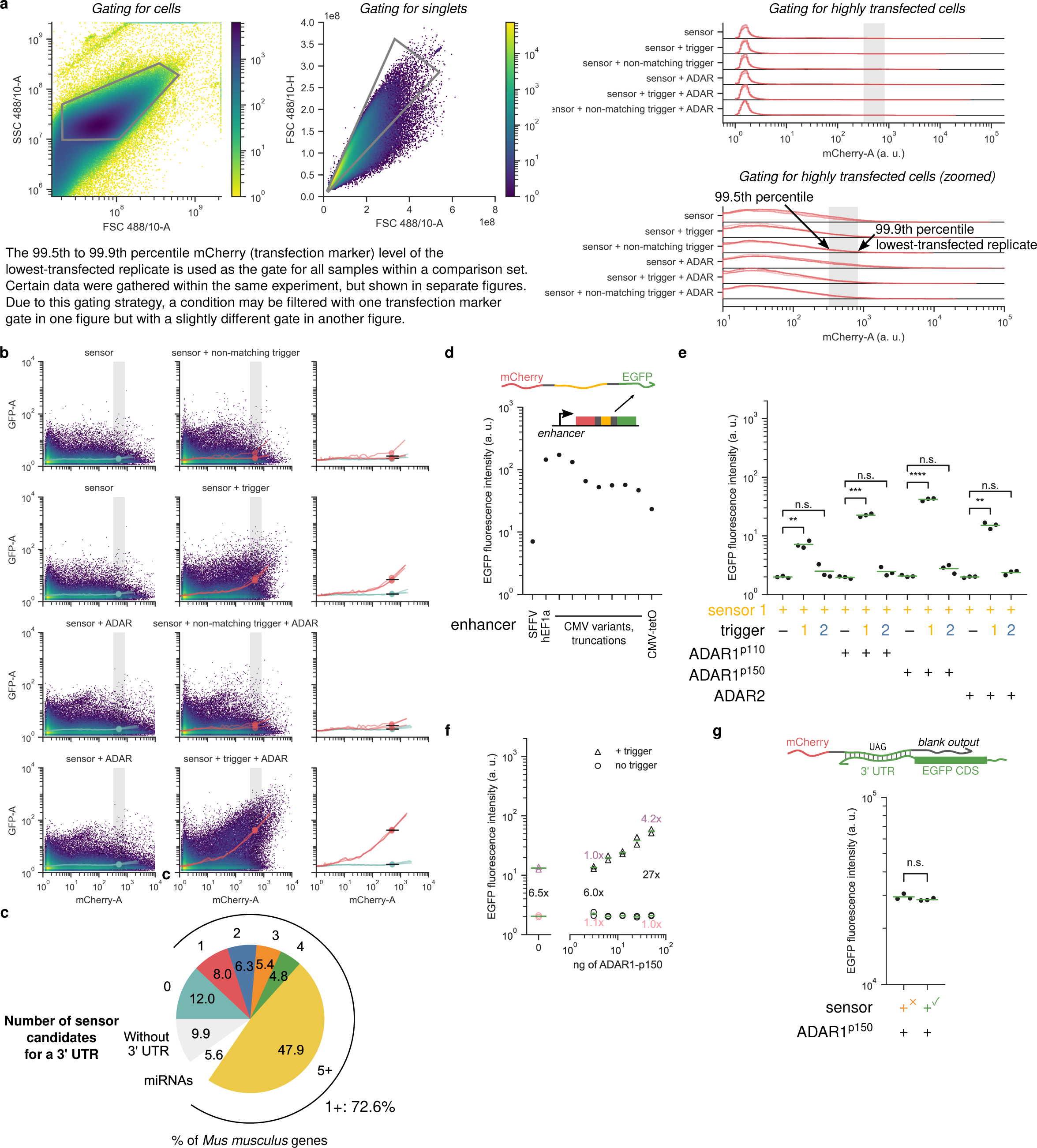
**a** Flow cytometry gating overview. **b** The ratio between the mean output fluorescence of the triggered and untriggered conditions depends on the chosen marker gate. Two-dimensional density plot showing the output (EGFP fluorescence, GFP-A) as a function of transfection marker levels (mCherry fluorescence, mCherry-A). The grey band indicates the chosen gate, as in subplot **a**. Traces indicate mean of EGFP fluorescence in small mCherry fluorescence bins. Points indicate mean in the chosen gate. Each replicate is a separate trace, but the 2D histogram combines all replicates. Rightmost column overlays all replicates. Black lines indicate the average of means across replicates, with the width indicating the gate. The means match those shown in main Figure 1c. **c** Mouse transcriptome analysis. **d** Enhancer choice affects baseline signaling, with SFFV giving the lowest baseline signal (EGFP fluorescence in the absence of trigger). Promoter variants were combined with the sensor using overlap extension PCR and transfected as linear fragments. **e** The p150 isoform of ADAR1 was the best ADAR at improving the dynamic range of RADAR output. **f** ADAR levels modulate sensor output. Purple numbers indicate the fold difference of the triggered case without ADAR, pink numbers indicate the fold difference to the untriggered case without ADAR, and black numbers indicate the fold activation upon adding trigger. **g** 3’ UTR sensor has marginal effect on trigger protein expression (EGFP fluorescence).

**Supplemental Figure 2.**
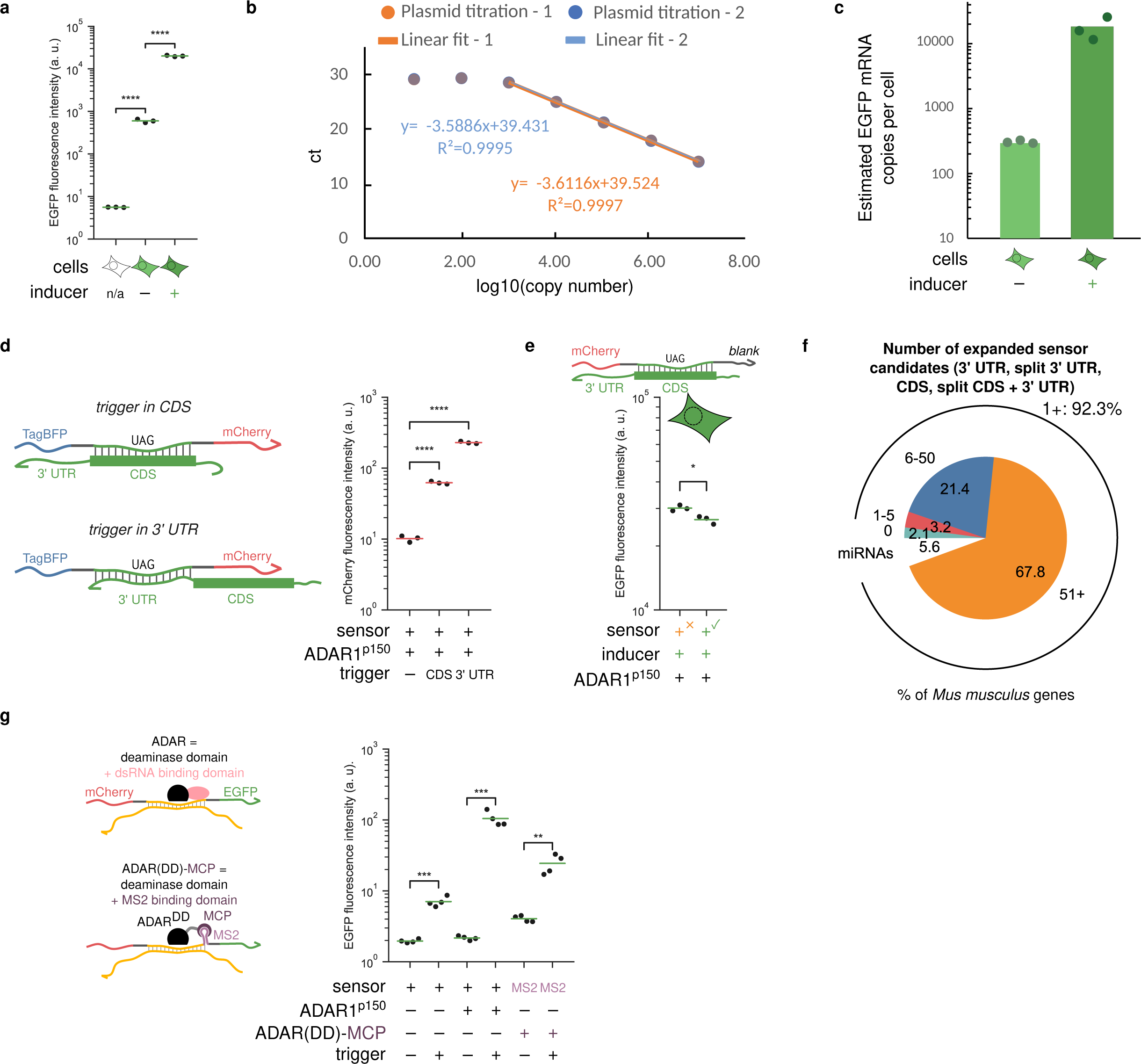
**a** The inducible trigger shows imperfect repression, as EGFP is detected even when inducer is not present. Parental (no EGFP) or inducible-EGFP-integrated cells were transfected with an unrelated sensor (TagBFP transfection marker, mCherry output) and ADAR1-p150 to measure the average EGFP expression. **b** Calibration curve with varying amounts of plasmid DNA of construct used to generate the inducible-EGFP-integrated cell line. Orange and blue (two replicates) overlapping appears grey in the plot. **c** Estimated number of the average mRNA molecules per cell in the inducible-EGFP-integrated cells, either with or without inducer. **d** A trigger sequence in the CDS performs more poorly compared to the same exact sequence placed into the 3’ UTR. **e** The sensor for a CDS sequence has marginal effect on the trigger mRNA protein expression (EGFP fluorescence reduced 1.14-fold). **f** Mouse transcriptome analysis using expanded set of trigger-sensor designs. **g** MS2 sequences directly downstream of the sensor sequence can be used to recruit ADAR(DD)-MCP.

